# Quantum and Classical Graph Convolutional Neural Networks for Protein Ligand Dissociation Constant Prediction

**DOI:** 10.1101/2025.11.20.689635

**Authors:** Azamat Salamatov, Jun Bai, Gowtham Atluri, Chaowen Guan

## Abstract

How long a drug stays bound to its target - the residence time - is now recognized as a stronger in vivo efficacy driver than binding affinity alone. Yet current machine learning (ML) models for dissociation kinetics (koff) ignore two critical sources of structure information: (1) the change in protein–ligand geometry across time, and (2) the ability to represent complex spatial interactions with fewer parameters. We extend a state-of -the -art spatial GNN for protein–ligand complexes with two innovations. First, we introduce a 2–timestep GCN+GRU model that learns structure changes before and after short molecular dynamics simulations. Second, we compress the model head using variational quantum circuits, preserving expressivity while removing 66% of parameters. On the full PDBbind-koff-2020 benchmark, temporal integration improves predictive accuracy, and quantum compression matches the full classical model’s accuracy at a fraction of its size. Our results show that time and quantum structure are underexplored, high-leverage axes for advancing kinetic ML models used in drug design.

## I. Introduction

In structure-based drug design, understanding the dynamics of protein-drug interactions is crucial, particularly from both thermodynamic and kinetic perspectives. Thermodynamics (e.g., equilibrium constants: *K*_*i*_, *K*_*d*_, *IC*_50_, etc.) describes the stability of the drug-protein complex at equilibrium, while kinetics (e.g., association and dissociation rates: *k*_on_, *k*_off_^1^) provides insights into how quickly the drug binds and unbinds from the target [23], [24]. Even drugs with similar binding strengths can differ significantly in their kinetic behaviors, which can influence their therapeutic effects [25]. Drugs that dissociate quickly generally require higher doses due to shorter action times, whereas those with slower dissociation rates stay bound longer, allowing for lower doses, enhanced selectivity, and potentially fewer side effects. These kinetic properties can therefore offer valuable guidance in optimizing drug efficacy and safety [25], [26], [27].

Currently, most studies assess the strength of drugtarget protein interactions using thermodynamic properties, such as the equilibrium binding constant (*K*_*i*_). These measurements are typically obtained from *in vitro* binding assays conducted in closed systems, where drug and protein concentrations remain constant. However, this setup does not reflect the dynamic, constantly changing conditions within the body. In the open, *in vivo* environment, drugs that bind or dissociate slowly may not reach equilibrium within the standard experimental timeframe, making it challenging to accurately capture their interaction characteristics under physiological conditions [23], [27].

Kinetic data on drug-target protein interactions are increasingly recognized as a more accurate indicator of a drug’s *in vivo* efficacy compared to thermodynamic data [26]. Recent studies have shown that the pharmacological activity of drugs in living systems is more strongly correlated with kinetic properties, such as binding and dissociation rates, than with thermodynamic equilibrium constants [25]–[29]. For instance, research by the Heitman group demonstrated that the intracellular efficacy of A2A adenosine receptor agonists correlates closely with their residence time (with an *R*^2^ of 0.90), while showing only a weak correlation with the equilibrium inhibition constant *K*_*i*_ (with an *R*^2^ of 0.13) [28]. This highlights the importance of considering binding kinetics, alongside affinity, to enhance drug development outcomes and improve the likelihood of therapeutic success [27].

## II. Related Work

Researchers are increasingly focused on creating computational approaches to predict dissociation rate constants (*k*_off_), aiming to complement experimental methods. Traditional approaches often rely on computationally intensive kinetic simulations to study drug dissociation pathways [30]. Recently, some studies have turned to quantitative structure-activity relationship (QSAR) models to improve predictive performance for dissociation kinetics [31], [32].

Several QSKR models have been reported in the literature, typically focused on specific drug targets. In 2016, Mei Hu’s group at Chongqing University developed a QSKR model for 37 HIV-1 protein inhibitors, using the 3D molecular force field Volsurf descriptor along with various physicochemical properties [31]. They employed partial least squares (PLS) and support vector machine (SVM) algorithms, achieving a correlation coefficient (*R*) of 0.772 on the test set for predicting dissociation rate constants. Similarly, in 2018, Wade et al. applied COMBINE analysis and PLS to construct a QSKR model for 66 HSP90 protein inhibitors, with a correlation coefficient (*R*^2^) of 0.86 on the test set [32]. These studies, along with others [5]–[14], have shown that QSKR models are feasible, although their limited dataset sizes and single-target focus restrict their broader applicability in drug development.

In 2020, Su et al. developed a more general-purpose QSKR model using a random forest algorithm on the newly curated PDBbind-*k*_off_-2020 dataset [34]. Their model, trained on 406 protein-ligand complexes, achieved a correlation coefficient (*R*) of 0.623 on the test set, demonstrating the potential of structure-based machine learning models for predicting *k*_off_.

That same year, Amangeldiuly et al. introduced a QSKR model using structural descriptors, specifically atom-type contact counts within a binding site radius, to predict unbinding kinetics [27]. This model was trained on a dataset of 501 protein-ligand complexes with experimentally measured *k*_off_ values. Using random forest algorithms, they achieved a correlation coefficient (*R*^2^) of 0.64, comparable to MD-based methods but with significantly reduced computational demands.

In 2023, Zhao et al. developed machine learning models for predicting *k*_off_ values, employing eight algorithms, including Bayesian Neural Network (BNN), random forest (RF), support vector machine (SVM), and extreme gradient boosting (XGBoost) [36]. Using descriptors based on van der Waals and electrostatic interaction energies, they conducted case studies on HSP90 and RIP1 kinase inhibitors, achieving state-of-the-art accuracy with the BNN model, which obtained correlation coefficients of *R*^2^ = 0.947 and *R*^2^ = 0.745 for the HSP90 and RIP1 datasets, respectively. This study demonstrates the effectiveness of machine learning, particularly BNN, for *k*_off_ prediction, suggesting these models are well-suited for high-throughput screening.

In 2024, Akhunzada et al. proposed a machine learning approach focusing on dynamic interaction fingerprints (IFPs) along the ligand unbinding pathway [35]. Using a dataset of 85 protein-ligand complexes specific to heat shock protein (HSP90) from the PDBbind-*k*_off_-2020 database, they generated dynamic IFPs via steered molecular dynamics (SMD) simulations. By incorporating these dynamic IFPs into a random forest (RF) regression model, they achieved a correlation coefficient (*R*^2^) of 0.80, outperforming static IFP-based models with an *R*^2^ of 0.60, thereby highlighting the value of dynamic interaction data in capturing unbinding kinetics.

### Our Contribution

While previous studies have advanced the modeling of *k*_off_ values, challenges persist in capturing the mutual information between proteins and ligands and in identifying key atoms and residues involved in these interactions. To address these challenges, we introduce two novel approaches for predicting the ligand dissociation constant (*k*_off_): a classical Graph Convolutional Network (GCN) and a quantum-enhanced GCN (QGCN). To our knowledge, this is the first application of both graph neural networks and quantum neural networks for predicting the ligand dissociation constant (*k*_off_). Our models utilize spatial data from 3D structural representations of protein-ligand complexes in the PDBbind-*k*_off_-2020 dataset [34], represented as graphs, to capture intricate interaction patterns. Additionally, we incorporate a GCN architecture adapted from [22], optimized using AutoDock Vina [33] distance terms, and integrate a bitransport information mechanism with physics-based distance terms to enhance prediction accuracy. Unlike other methods, our approach not only effectively captures mutual information between proteinligand pairs but also highlights essential atoms in the ligands and residues in the proteins. These innovations present a promising framework for advancing structure-based drug design.

## III Dataset

### A. Dataset Collection and Data Mining Approach

In this study, the PDBbind-*k*_off_-2020 dataset [34] was used to benchmark protein–ligand dissociation rate constant (*k*_off_) prediction. This dataset is a collection of protein–ligand complexes for which experimentally measured *k*_off_ values are available. To prepare the dataset, we carefully filtered the protein-ligand pairs based on the following criteria: (i) all measurements had to be specific values rather than approximations or ranges; (ii) ligand structures had to match between the PDBbind-*k*_off_-2020 dataset and the chemical component dictionaries in the PDB databank; (iii) proteins were required to have only one C*α* atom per residue, (iv) it was possible to read and parse the files. Initially, the dataset contained 680 protein-ligand complexes, from which 423 were retained. After filtering, we split the dataset into training, validation, and test sets in 70:15:15 ratio, respectively.

3D spatial features of an individual protein and ligand were read and parsed from PDB and MOL2 files, respectively [4], [5], [18].

### B. Feature Representation

#### 1) Protein Representation

##### Node Feature Matrix

In our model, protein 3D structures are represented as graphs, where nodes correspond to 20 different amino acid residues, and edges connect nonconsecutive amino acid residues whose C*α* atoms are within an 8 Å distance. Each node in this graph has a 41-dimensional feature vector that combines primary sequence information, normalized position, and evolutionary relationships:

(1) *Primary sequence information;*A 20-dimensional one-hot encoding is used to represent the amino acid types in the protein sequence; (2) *Normalized position*: The position of each amino acid in the sequence is scaled by the total sequence length (number of residues in this sequence); (3) *Evolutionary relationships*: Using the BLOSUM62 matrix [37], which captures evolutionary similarities among amino acids, a 20-dimensional vector from the matrix represents each amino acid type.

Together, these features yield a 41-dimensional vector for each node in the protein graph.

##### Adjacency Matrix

The adjacency matrix represents connections between amino acid residues as nodes, with edges indicating non-covalent interactions between these residues. An edge is created if the Euclidean distance between the *C*_*α*_ atoms of two amino acids is less than 8 *Å*. This distance map is converted into an adjacency matrix by assigning a value of 1 for edges (distances ≤ 8 *Å*) and 0 otherwise.

##### 2) Ligand Representation

In the ligand graph, nodes represent atoms, and edges denote chemical bonds, generated using RDKit [38]. The initial node features for ligands include: (1) *Atom type*: 10-dimensional one-hot encoding of atom types (H, C, N, O, F, P, S, Cl, Br, I), only these 10 atoms appear in the whole dataset; *Atom degree*: 5-dimensional one-hot encoding based on atom connectivity; (3) *Atom explicit valence*: 7-dimensional one-hot encoding for the explicit valence of each atom; (4) *Atom implicit valence*: 2-dimensional one-hot encoding for the implicit valence; (5) *Aromaticity*: A 2-dimensional feature indicating whether the atom is aromatic.

These features result in an 26-dimensional vector per atom. For edges, a 6-dimensional feature vector represents six bond types, including single, double, triple, aromatic bonds, as well as conjugated and ring bonds.

##### 3) Interaction Representation

An interaction-based subgraph captures the pocket-ligand interaction features. Nodes in this subgraph include protein amino acid residues and ligand atoms: - Amino acid residues nodes: Each node has a 41-dimensional feature vector, as defined in the protein representation. - Ligand atom nodes: Each node is represented by the 26-dimensional feature vector described in the ligand representation.

Edges in this subgraph are created if the Euclidean distance between a heavy atom from the ligand and an amino acid C*α* atom is less than 8 Å. The edge feature is defined as 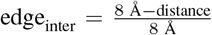 for distances under 8 Å, and is set to 0 otherwise.

### C. Vina Distance Optimization Terms

In our study, the Vina distance optimization terms incorporate the same weighted scoring terms as used in AutoDock Vina [33]. These terms are composed of five conformation-dependent terms and one conformation-independent term. The five conformation-dependent terms represent intermolecular contributions, including three terms for steric interactions, one term for hydrophobic bonding, and one for hydrogen bonding. The detailed setup of Vina terms is available in Appendix B.

All code implementation were done on Python programming language. PyTorch deep learning library [41], and Deep Graph Library (DGL) [42] were used to construct graphs, and train the Graph Convolutional Networks (GCN). PennyLane [44] and Qiskit [40] quantum machine learning libraries were used to construct quantum neural networks [43].

## IV Quantum Data Encoding

In quantum computing, data from classical systems must be encoded into quantum states. This section discusses two encoding schemes used in our work: *Amplitude Encoding* and *Variational Encoding*. We demonstrate each encoding with an example. Note: the basics of quantum computing are covered in Appendix A.

### A. Amplitude Encoding

Amplitude encoding maps a classical data vector into the amplitudes of a quantum state. The visual representation of the quantum circuit for amplitude encoding is provided in Fig. 15. Given a normalized classical vector **x** = [*x*_0_, *x*_1_, …, *x*_*N−*1_], it is encoded as a quantum state:

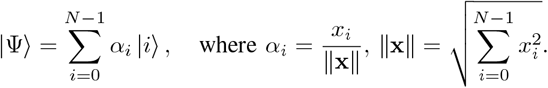

This encoding reduces the number of qubits needed for large datasets, as an *N* -dimensional vector requires only log_2_(*N*) qubits.

### B. Variational Encoding

Variational encoding maps classical data into quantum states by using the values in the data array as rotation angles for quantum gates, see Fig. 16 for the visual representation of the quantum circuit. For a given classical vector **x** = [*x*_1_, *x*_2_, …, *x*_*n*_], the encoding uses parameterized single-qubit gates, such as *R*_*y*_ and *R*_*z*_, to prepare the quantum state:

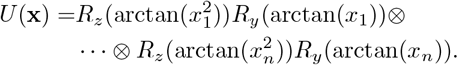

## V. Model Construction

In this work, we employed (1) a graph convolutional neural network (GCN) architecture from [22] incorporating Vina distance optimization terms to predict proteinligand dissociation constant (*k*_off_), (2) a concatenation of two GCNs with described architecture processed by a Gated Recurrent Unit (GRU) layer and trained on protein-ligand structures before and after 2-ns Molecular Dynamics (MD) simulation, (3) a Quantum GCN with reduced number of parameters and augmented by Quantum Layer - Variation Quantum Circuit.

### A. GCN Architecture

As illustrated in Fig. 2, the architecture begins with embedding layers that convert node and edge vectors into fixed-dimensional representations. The model is composed of three primary GNN blocks designed to learn features from protein structures, ligand graphs, and pocket-ligand interaction structures, respectively. The details of the GNN setup are as follows.

**Fig. 1.**
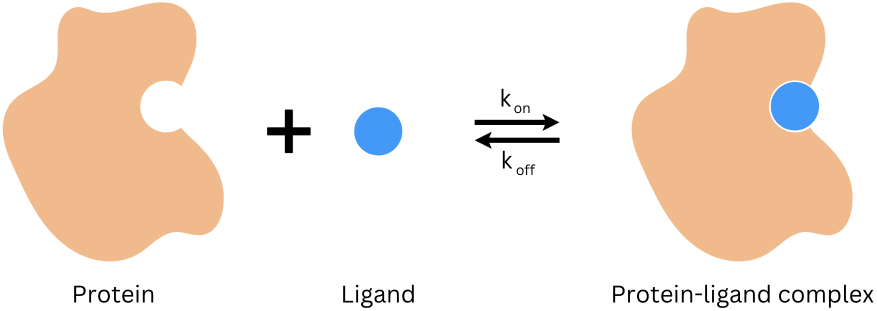
Protein-ligand binding reaction pathway with the association rate constant (*k*_on_) and the dissociation rate constant (*k*_off_)

**Fig. 2.**
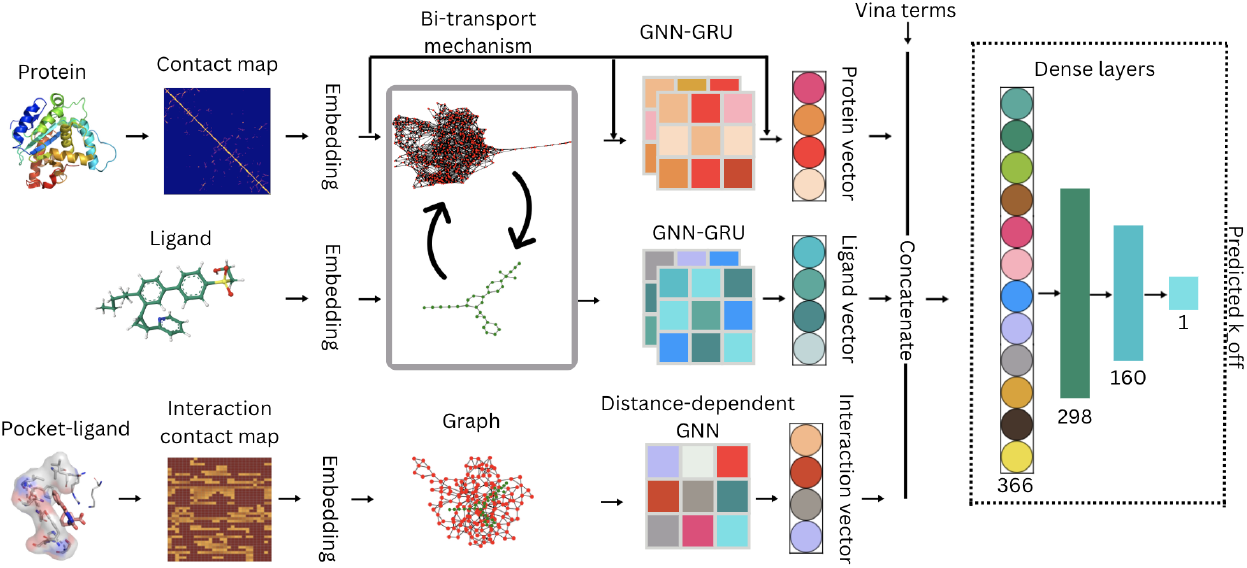
Three main GNN blocks were developed to extract features from the protein structure, ligand structure, and pocket-ligand interaction structure, as proposed in GraphscoreDTA [22]. In the protein and ligand blocks, a bitransport information mechanism was introduced to facilitate communication between the protein and ligand representations. Separate GNNs were then employed to capture structural features of the protein and ligand, integrating multihead attention, GRUs, and skip connections within the networks. For the pocket-ligand interaction block, distance-dependent GNNs were used to model the pocket-ligand structure. Finally, the features learned from all three blocks, along with Vina distance optimization terms, were combined and passed through three dense layers to predict binding affinity.

First, a bitransport information mechanism was introduced to facilitate information exchange between the protein and ligand representations. Then, the amino acid and atom features were processed through two iterations of protein and compound GNNs, producing updated feature representations. In the second iteration, a skip connection was added between the embedding vector and the GNN input, followed by another skip connection linking the embedding vector to the GNN output. In the interaction graph, the amino acid and atom features were processed by a single GNN layer to update the interaction features, and a skip connection was applied between the embedding vector and the GNN output.

Next, a multihead attention mechanism was incorporated into both the protein and ligand GNNs to adjust the importance of individual atoms within the ligand and specific residues within the protein. Additionally, gated recurrent units (GRUs) were employed to control the information flow between the general node and the super node within the ligand and protein graphs. To enhance predictive performance, Vina distance terms, capturing intermolecular interactions and ligand flexibility, were included in the model. As discussed in III-C, these terms were derived from AutoDock Vina, a widely used tool for molecular docking and virtual screening, and include three steric interaction terms (gauss1, gauss2, and repulsion), a hydrophobic interaction term, and a hydrogen bond term. Ligand flexibility is represented by the number of rotatable bonds among the ligand’s heavy atoms.

Finally, the features generated by the three GNN blocks, along with the Vina distance optimization terms, were concatenated and passed through three dense layers. The first two dense layers consist of 390 and 320 nodes, each followed by a PReLU activation function, while the third layer has 160 nodes and leads to the output. To fine-tune the model, the validation set was used for parameter tuning, and the model was trained for 100 epochs. The version of the model achieving the best validation performance was selected. The chosen hyperparameters are detailed in Table I.

**TABLE I.** Hyperparameters for Different Model Blocks.

| Blocks | Hyperparameters | Settings for GCN | Settings of Quantum GCN |
| --- | --- | --- | --- |
|  | Epoch | 80 | 50 |
|  | Batch size | 8 | 4 |
|  | Optimizer | AdamW | AdamW, RMSProp |
|  | Learning rate | 1.2e-4 | 1.2e-3 |
|  | Weight decay | 1e-6 | 1e-5 |
| Ligand GNN block | Iteration | 2 | 2 |
|  | Graph node feature size | 120 | 40 |
|  | Hidden size (node) | 120 | 40 |
|  | Graph edge feature size | 8 | 6 |
|  | Attention heads | 4 | 4 |
|  | Activation function | LeakyReLU, Tanh, Sigmoid | LeakyReLU, Tanh, Sigmoid |
| Protein GNN block | Iteration | 2 | 2 |
|  | Graph node feature size | 120 | 40 |
|  | Hidden size (node) | 120 | 40 |
|  | Attention heads | 4 | 4 |
|  | Activation function | LeakyReLU, Tanh, Sigmoid | LeakyReLU, Tanh, Sigmoid |
| Pocket-ligand interaction distance-based GNN block | Iteration | 1 | 1 |
|  | Graph node feature size | 120 | 40 |
|  | Distance cutoff | 8 Å | 8 Å |
|  | Depth | 3 | 3 |
|  | Activation function | ReLU | ReLU |
| Fully connected neural network block | Number of neurons | [366, 298, 160] | [126, 96, 32] |
|  | Activation function | PReLU | PReLU |

#### 1) Bitransport information mechanism

In this study, we utilized a bitransport information mechanism (introduced in [22], here adapted by us for *k*_off_ prediction) for information exchange between protein and ligand feature representations. This mechanism relies on multihead attention and position-aware attention to capture the distinct contributions of each component.

For the ligand feature updates, multihead attention was applied to transform the protein feature matrix *X*_*p*_ ∈ ℝ^*n*×*c*^ into a unified global descriptor *G*_*p*_ ∈ ℝ^1×*c*^. Here, we used *h* = 8 attention heads to aggregate information effectively. The global representation *G*_*p*_ for the protein is defined as:

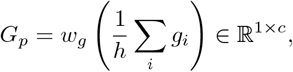

where *w*_*g*_ is a learnable parameter, *c* represents the feature dimension, and *g*_*i*_ denotes each attention head output, defined by:

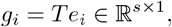

with *T* derived by applying a transformation on *X*_*p*_ as follows:

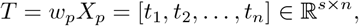

where *w*_*p*_ is another learnable parameter. Additionally, multihead attention employs a matrix *B*, constructed by:

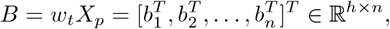

where each attention score 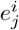 for position *j* is given by:

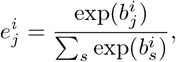

where *s* represents the feature dimension, and *n* is the total number of amino acids in the protein.

Position-aware attention then updates each ligand atom’s features. The initial ligand features *X*_*l*_ ∈ ℝ^*m*×*c*^ are modified as follows:

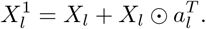

Here, position-aware attention for ligand features is computed by:

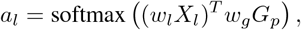

where *w*_*l*_ and *w*_*g*_ are learnable parameters, and *m* is the number of ligand atoms. This approach is repeated in the protein to get updated amino acid residue features.

#### 2) Multihead Attention Mechanism

In the ligand atom graph, in addition to the standard atom nodes, a virtual “super node” was introduced to gather information from the actual atom nodes. This multihead attention mechanism was applied to assess the contribution of each individual atom node. For transferring information from general atom nodes to the super node, a *K*-head attention mechanism was implemented as follows:

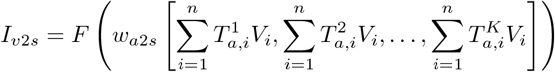

where *F* denotes the tanh activation function, *n* is the number of ligand atoms, and *w*_*a*2*s*_ represents a learnable weight. In this implementation, *K* = 4 attention heads were used.Each 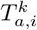 is computed using a softmax function, which assigns weights based on the specific attention parameters:

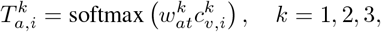

where *i* represents the index of ligand atoms, and 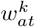 is a learnable parameter.

The values 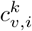 are calculated by combining features from the atom nodes and the super node using the following formula:

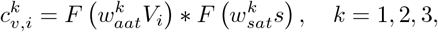

where 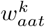 and 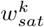 are learnable weights, *V*_*i*_ represents the atom node features, and *s* represents the features of the super node.

This process is similarly applied in the protein graph, where information is transferred and updated using the same multihead attention approach.

Here is a paraphrased version of the text with mathematical notation included:

#### 3) Gated Recurrent Units

To model dependencies across different time scales, a Gated Recurrent Unit (GRU) [45] was incorporated into the architecture. This GRU mechanism helps capture the temporal dependencies among nodes in both individual atom and super nodes. Two GRUs were employed to manage the proportion of information shared between individual atoms and the super node.

The feature update for the super node can be defined as:

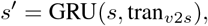

where tran_*v*2*s*_ is the transformed information from atom nodes to the super node, computed as:

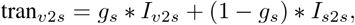

with *g*_*s*_ being the gating mechanism determined by:

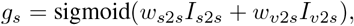

where *w*_*s*2*s*_ and *w*_*v*2*s*_ are learnable parameters.

The update of features for each individual atom node *V*_*i*_ is calculated as:

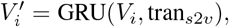

where tran_*s*2*v*_ is the information transferred from the super node back to the individual atom nodes, defined by:

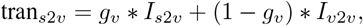

and the gating mechanism *g*_*v*_ is calculated as:

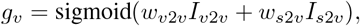

where *w*_*v*2*v*_ and *w*_*s*2*v*_ are learnable weights.

In this setup, tran_*v*2*s*_ denotes the transformed information moving from atom nodes to the super node, while tran_*s*2*v*_ represents the flow of processed information from the super node back to the atom nodes. The terms *I*_*v*2*s*_, *I*_*s*2*s*_, *I*_*v*2*v*_, and *I*_*s*2*v*_ refer to different types of interaction information between nodes.

#### 4) Graph Neural Network

The graph neural network (GNN) was developed to capture the ligand graph features, incorporating two types of nodes—the individual atom nodes and a virtual super node—as well as a single type of edge encoding. Five distinct types of information were utilized to compute and update the node features within this graph: (i) information from neighboring atoms; (ii) data from the super node; (iii) transformed data transferred from individual atoms to the super node; (iv) transformed information passed from the super node back to individual atoms; and (v) edge-related information. The update process for the super node features was implemented using the tanh activation function (see Fig. 6-A).

**Fig. 3.**
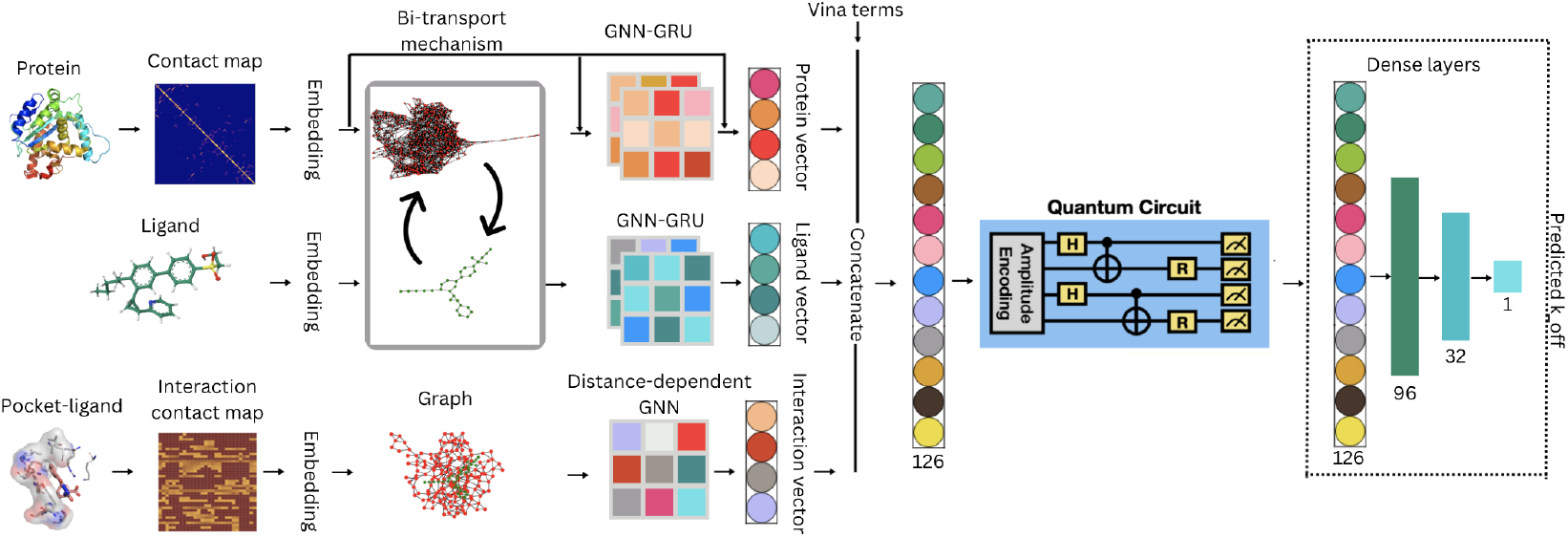
The Quantum GCN architecture. The output vector of size 126 is trained on the Variational Quantum Circuit and followed by classical post-processing step: fully connected dense layers. The quantum circuit in this figure is simplified for demonstration purposes. The complete variation quantum circuits are provided in Fig. 15 and Fig. 16.

**Fig. 4.**
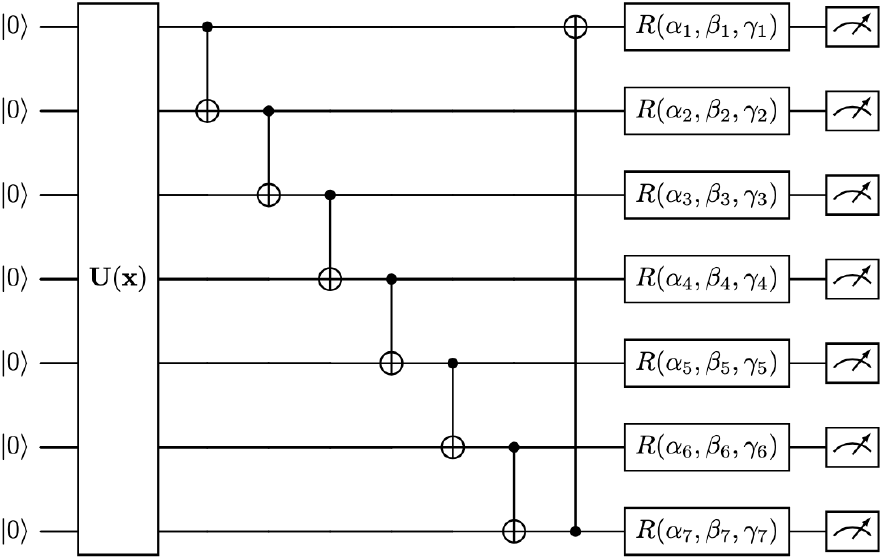
Variational quantum circuit architecture for the classifier with amplitude encoding. The initial VQC block encodes the vector produced by the graph convolution operation. An *N* -dimensional vector is transformed into a quantum state using log_2_(*N*) qubits. In this study, *N* = 10. The *U* (**x**) function represents the quantum routine for amplitude encoding. Trainable parameters, denoted by *α*_*i*_, *β*_*i*_, and *γ*_*i*_, are optimized during training.

**Fig. 5.**
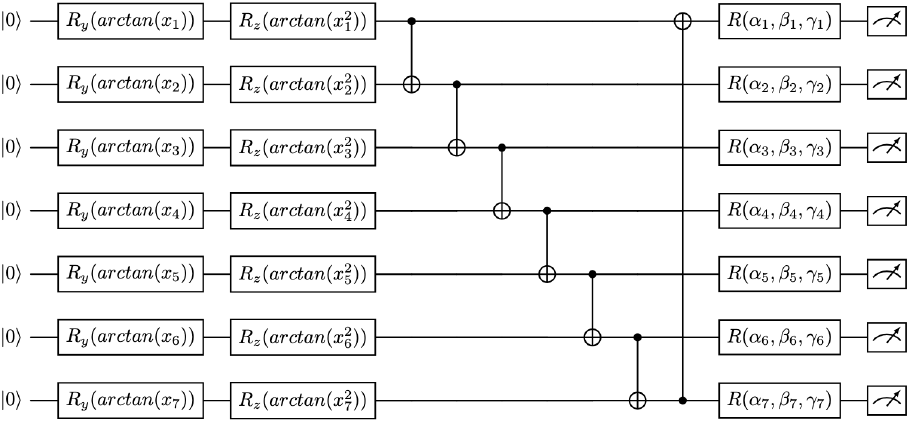
Variational quantum circuit block with variational encoding. This encoding block employs *R*_*y*_ and *R*_*z*_ rotation gates, where the rotation angles are parameterized by arctan(*x*_*i*_) and arctan 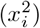 for each qubit. The parameters *α*_*i*_, *β*_*i*_, and *γ*_*i*_ are trainable and optimized during the learning process.

**Fig. 6.**
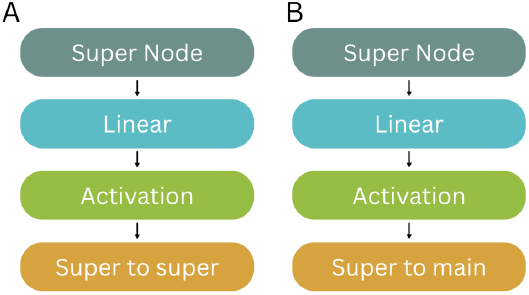
Information transfer of super node

To transfer information from individual atoms to the super node, a multihead attention mechanism was employed (as shown Fig. 7). Similarly, information from the super node back to individual atoms was updated using the tanh activation function (see Fig. 6-B). The aggregation of information from neighboring atoms was achieved through one-step neighbor information collection along graph edges (see Fig. 8).

**Fig. 7.**
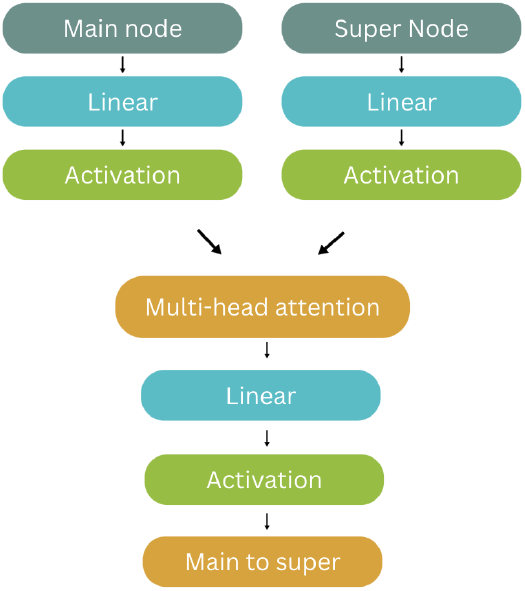
Information transfer from main node to super node

**Fig. 8.**
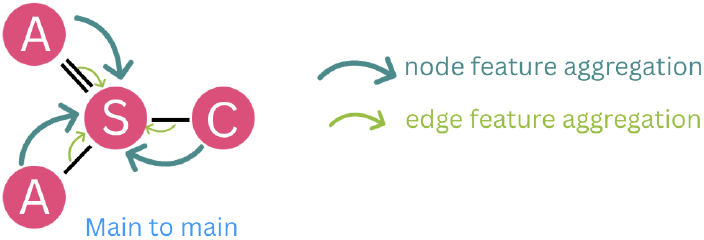
Atom information gathered from its neighbor atoms

To refine the feature representations, two GRUs were applied separately to update features of individual atoms and the super node. This GNN structure iterated twice to update the ligand node features fully. The protein graph features were updated through a similar GNN process, differing only in that edge features were excluded from the graph updates.

#### 5) Pocket–Ligand Distance-Dependent Graph Neural Network

In the pocket-ligand interaction graph, a radial pooling mechanism based on distance was employed to capture the interaction edges. Each interaction edge is defined as:

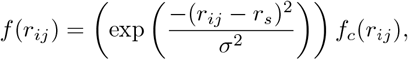

where *r*_*ij*_ represents the interaction distance between a protein residue and a ligand atom, and *r*_*s*_ and *σ* are learnable parameters. The term *f*_*c*_(*r*_*ij*_) is a cutoff function, defined as:

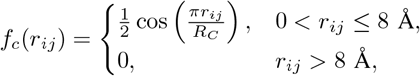

where *R*_*C*_ is set to 8 Å.

Following this, the graph node features are updated by aggregating information from neighboring nodes along different edges, as well as from self-node contributions. Ultimately, three iterations of distance-based radial pooling within the pocket-ligand interaction graph neural network are performed to enhance and update the node features.

### B. 2-GCN-GRU Architecture

In order to capture the temporal relationship between protein-ligand complexes we applied the described GCN to 2 datasets, where both of them have the same proteinligand complexes but have different 3D structures. The first dataset captures the initial structures of complexes (before the chemical reaction). The second dataset contains the structures of the same complexes after 2-ns Molecular Dynamics (MD) simulation (after the chemical reaction).

Both of these datasets were passed through two GCNs with same architecuture but distinct parameters. Their outputs of size 126 and 126 were concatenated into an array of size 252 and fed into a new Gated Recurrent Unit (GRU) layer. The introduced GRU layer is expected to capture the temporal relationship between the 2 timesteps of the complexes.

The output of the GRU is processed in a similar manner by passing through fully connected layers.

### C. Quantum GCN Architecture

The proposed Quantum GCN framework consists of three main components: (1) the described Graph Convolutional Network (GCN), (2) a Variational Quantum Circuit (VQC), and (3) a classical post-processing stage, see Fig 3. The idea of introducing the quantum layers - VQCs - is to reduce the number of parameters in the GCN, while preserving its accuracy using quantum advantage. Thus, we were able to reduce the complexity of the graph neural network, which had input size of 366, into a much smaller neural network of size 126. Refer to Table I for the details of GCN compression.

Initially, the original complex structures are processed through the reduced GCN, which outputs a feature vector of size 126. Then, the vector of size 126 is then padded with two additional zeros to reach a size of 128, aligning it with the requirement of 2^*n*^ for amplitude encoding.

Next, the padded vector is encoded into a quantum state using amplitude encoding (as described in Section A-A) to minimize the number of qubits required. The resulting quantum state is subsequently processed through a series of variational operations. Specifically, the architecture includes two VQCs separated by a tanh activation function. The first VQC performs both the amplitude encoding and variational transformations, Fig. 15. The Pauli-Z (*Z*-gate) expectation values from the first VQC are then re-encoded using the variational encoding scheme and passed to the second VQC for further variational transformations, Fig. 16.

Finally, the measured expectation values from the second VQC block are fed into a fully connected neural network, which processes the data to output the predicted *k*_off_ value.

#### 1) Optimization of Quantum Circuits

For gradient-based optimization, the parameter-shift rule [46], [47] is employed to compute the gradients of quantum functions. This approach has been highly effective in VQC-based quantum machine learning tasks [48]–[54]. For a quantum function *f* (*x*; *θ*) with an observable 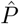, the gradient is calculated as:

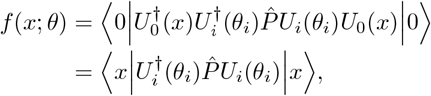

where *x* is the classical input from the GCN, *U*_0_(*x*) prepares the quantum state based on the classical data, *U*_*i*_(*θ*_*i*_) represents the single-qubit rotation generated by the Pauli operators *X, Y*, and *Z*, and *i* is the index of the circuit parameter being evaluated. The gradient of the function *f* with respect to the parameter *θ*_*i*_ is given by:

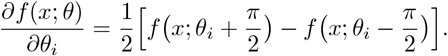

With the ability to calculate quantum gradients, various classical optimization algorithms can be applied. For this study, the RMSProp optimizer is used, which is a variant of gradient descent with an adaptive learning rate. The circuit parameters *θ* are updated as follows:

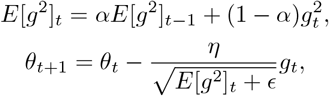

where *g*_*t*_ is the gradient at step *t* and *E*[*g*^2^]_*t*_ is the weighted moving average of squared gradients, with the initial value 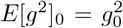. The hyperparameters are set as follows: learning rate *η* = 0.01, smoothing constant *α* = 0.99, and *ϵ* = 10^*−*8^.

## VI Evaluation

### A. Evaluation Metrics

We used the training and validation loss graphs to show the convergence of our models. Most of the approaches converge at epochs 40-50. For the demonstration of Quantum Advantage, we compared the loss of Quantum GCN which used the reduced number of parameters, with a similar reduced analog without the quantum layers. It is clearly seen that QGCN has lower loss (approx. 0.0023) compared to the reduced GCN with loss (approx. 0.0047)

**Fig. 9.**
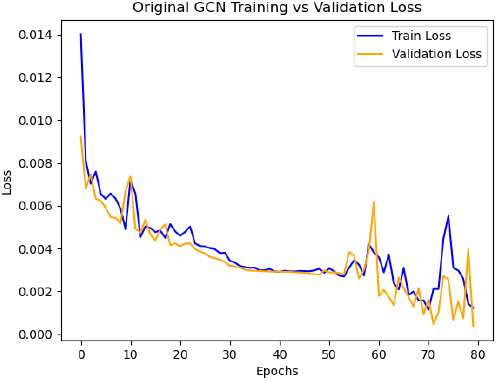
Original GCN Training vs Validation Loss

**Fig. 10.**
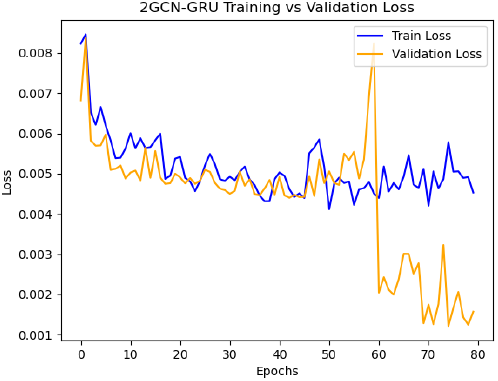
2GCN-GRU Training vs Validation Loss

**Fig. 11.**
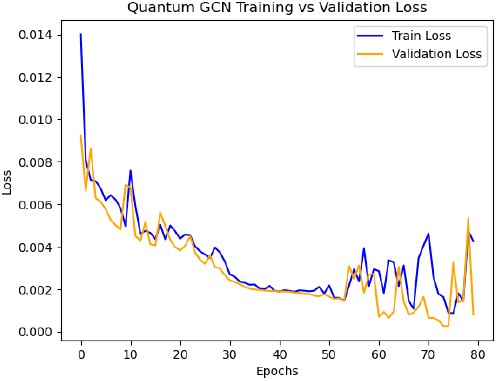
Quantum GCN Training vs Validation Loss

**Fig. 12.**
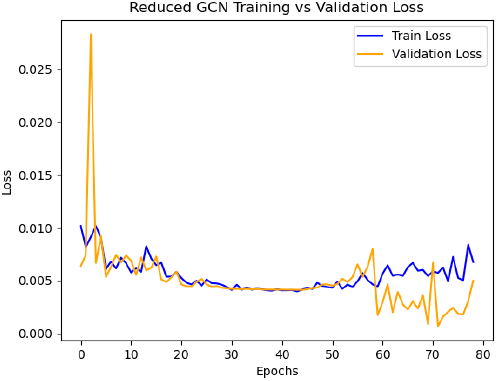
Reduced GCN Training vs Validation Loss

We used the following metrics to assess the predictive accuracy of our models: root mean squared error (RMSE) and mean absolute error (MAE). The formulas for each metric are as follows:

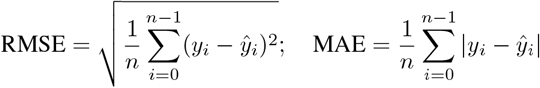

where *ŷ*_*i*_ is the predicted *k*_off_ value for the *i*-th sample, *y*_*i*_ is the actual *k*_off_ value, *n* represents the number of samples.

According to Table II the original GCN and Quantum GCN have comparable and even higher MAE and RMSE metrics for prediction *k*_off_ values. This indicates the supremacy of the Graph Neural Networks for spacial representation of proteins and ligand. In addition, the Quantum GCN even with lower number of parameters is approximately as precise as the original GCN initially proposed in [22], but upgraded to be used for *k*_off_ predition.

**TABLE II.** Performance comparison of various models based on RMSE and MAE. Bold values highlight the best performance in each metric.

| Author | RMSE | MAE |
| --- | --- | --- |
| GCN | **0.242** | **0.114** |
| GCN + 2 timestep GRU | 0.313 | 0.191 |
| Quantum GCN | **0.276** | **0.115** |
| Su et al. | 0.85 | — |
| Amangeldiuly et al. | 0.82 | 0.59 |
| Zhao et al. | **0.220** | 0.184 |
| Akhunzada et al. | 0.38 | 0.29 |

Although the Quantum GCN achieves comparable accuracy to the full classical GCN while using fewer parameters, our findings also highlight model-specific limitations. Despite the implementatoin of temporal structure, the 2GCN-GRU model, showed higher RMSE and MAE, suggesting that two timesteps of MD (0 ns and 2 ns) may not contain sufficiently informative structural changes for *k*_off_ prediction. The Quantum GCN, on the other hand, generalizes nearly as well as the high-capacity classical baseline, but still slightly underperforms the model of Zhao et al. in RMSE. This indicates that while quantum compression effectively reduces model size, further improvements in encoding schemes, circuit depth, and hybrid architectures may be needed to consistently outperform classical counterparts. Overall, these results provide evidence that spatial GNN representations are the dominant factor in *k*_off_ prediction, and that quantum layers can be integrated without degrading predictive accuracy.

## VII Conclusion and Future Discussion

In this work, we introduced three architectures for predicting protein–ligand dissociation constants (*k*_off_): a classical GCN model, a temporal 2GCN-GRU model, and a Quantum GCN that incorporates Variational Quantum Circuits to reduce the number of trainable parameters. Across the full PDBbind-koff-2020 benchmark, our results demonstrate that spatial GNNs remain highly effective for modeling protein–ligand interactions, outperforming previous QSKR approaches by a large margin. The Quantum GCN matches the accuracy of the full classical model while using a substantially smaller parameter set, highlighting the promise of quantum-based compression in structure-based learning.

Looking forward, several research directions emerge. First, temporal models may benefit from longer or better-selected MD trajectories, as two timesteps appear insufficient to capture relevant unbinding dynamics. Second, improving quantum encodings—such as data-reuploading, entangling layers, or QSVT-inspired blocks—may further enhance accuracy and robustness. Third, integrating physically motivated features such as path sampling metrics, interaction energies, or dynamic fingerprints could strengthen the model’s ability to capture long-timescale kinetic effects. Finally, scaling these models to larger datasets and benchmarking them on prospective, experimentally measured *k*_off_ values will be essential for assessing their translational impact in drug discovery.

## Code and Data Availability

The code used in this study is publicly available at https://github.com/asalamatov/Koff_Q_GCN. The dataset used in this study is also publicly available from the PDBbind database and can be accessed at https://www.pdbbind-plus.org.cn/.

## Footnotes

1 In the Fig. 1, the association rate constant *k*_on_ defines the speed at which a ligand binds to its target protein, often measured in units of M^−1^s^−1^. A high *k*_on_ value suggests a ligand rapidly associates with the receptor, which can be crucial for fast-acting drugs. Conversely, *k*_off_ represents the dissociation rate constant, or how quickly the ligand unbinds from the receptor, with units of s^−1^. A low *k*_off_ value indicates the ligand remains bound for a longer duration, contributing to prolonged drug efficacy.

