## Appendix for "Quantum and Classical Graph Convolutional Neural Networks for Protein Ligand Dissociation Constant Prediction"

### APPENDIX A
### BASICS OF QUANTUM COMPUTING

Quantum computing operates on principles fundamentally different from classical computing. The basic unit of quantum information is the *quantum bit*, or *qubit*, which exists in a two-dimensional Hilbert space. In this section, we provide a foundational overview of quantum states, vector representations, norms, and amplitudes.

***D1. Quantum States:*** A qubit is represented as a *quantum state vector* in the Hilbert space $\mathbb{C}^2$, with basis states $|0\rangle$ and $|1\rangle$. These basis states correspond to the classical binary states 0 and 1 and are defined as:

$$|0\rangle = \begin{pmatrix}1\\0\end{pmatrix}, \quad |1\rangle = \begin{pmatrix}0\\1\end{pmatrix}.$$

The general state of a qubit, denoted by $|\psi\rangle$, is a *linear combination* (or superposition) of these basis states:

$$|\psi\rangle = \alpha\,|0\rangle + \beta\,|1\rangle\,,$$

where $\alpha, \beta \in \mathbb{C}$ are complex numbers called *amplitudes*. The squared magnitudes of these amplitudes, $|\alpha|^2$ and $|\beta|^2$, represent the probabilities of measuring the qubit in the $|0\rangle$ and $|1\rangle$ states, respectively. These probabilities must sum to 1, leading to the normalization condition:

$$|\alpha|^2 + |\beta|^2 = 1.$$

***D2. The Bloch Sphere:*** The state of a single qubit can also be visualized as a point on the surface of a three-dimensional sphere, known as the *Bloch Sphere*. This representation provides a geometric interpretation of the quantum state. The state $|\psi\rangle$ can be parameterized as:

$$|\psi\rangle = \cos\left(\frac{\theta}{2}\right)|0\rangle + e^{i\phi}\sin\left(\frac{\theta}{2}\right)|1\rangle\,,$$

where $\theta \in [0, \pi]$ and $\phi \in [0, 2\pi]$ are spherical coordinates. Here:

- The north pole of the sphere represents the state $|0\rangle$.
- The south pole represents the state $|1\rangle$.
- Points on the surface represent superpositions of $|0\rangle$ and $|1\rangle$.

The Bloch Sphere is shown below:

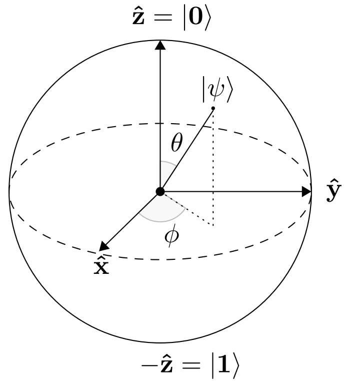


Fig. 13. The Bloch Sphere representation of a single qubit. The state $|\psi\rangle$ is parameterized by angles $\theta$ and $\phi$.

This representation emphasizes that the state of a qubit is not restricted to binary values but can exist in a continuum of superpositions.

***D3. Multi-Qubit Systems:*** For $n$ qubits, the quantum state resides in a $2^n$-dimensional Hilbert space. The basis states for two qubits are:

$$|00\rangle\,, |01\rangle\,, |10\rangle\,, |11\rangle\,,$$

represented as column vectors:

$$|00\rangle = \begin{pmatrix}1\\0\\0\\0\end{pmatrix}, \quad |01\rangle = \begin{pmatrix}0\\1\\0\\0\end{pmatrix}, \quad |10\rangle = \begin{pmatrix}0\\0\\1\\0\end{pmatrix}, \quad |11\rangle = \begin{pmatrix}0\\0\\0\\1\end{pmatrix}.$$

The general state of a two-qubit system is:

$$|\Psi\rangle = \alpha_0 |00\rangle + \alpha_1 |01\rangle + \alpha_2 |10\rangle + \alpha_3 |11\rangle,$$

where $\sum_{i=0}^{3} |\alpha_i|^2 = 1$.

***D4. Norms and Normalization:*** The *norm* of a quantum state vector quantifies its length. For a state $|\psi\rangle$, the norm is defined as:

$$\|\psi\| = \sqrt{|\alpha|^2 + |\beta|^2}.$$

In quantum mechanics, state vectors must always be normalized, meaning their norm equals 1. If a vector is not normalized, it can be rescaled by dividing each amplitude by the norm:

$$|\psi_{\text{normalized}}\rangle = \frac{|\psi\rangle}{\|\psi\|}.$$

***D5. Measurement and Probabilities:*** When a quantum state $|\psi\rangle = \alpha |0\rangle + \beta |1\rangle$ is measured in the computational basis $\{|0\rangle, |1\rangle\}$, the outcome is probabilistic:

- The probability of measuring $|0\rangle$ is $|\alpha|^2$,
- The probability of measuring $|1\rangle$ is $|\beta|^2$.

The measurement collapses the state into the observed basis state, and the superposition is lost.

***D6. Tensor Product for Multi-Qubit States:*** To combine multiple qubits, the *tensor product* is used. For example, given two qubits $|\psi\rangle_1 = \alpha_1 |0\rangle + \beta_1 |1\rangle$ and $|\psi\rangle_2 = \alpha_2 |0\rangle + \beta_2 |1\rangle$, their combined two-qubit state is:

$$|\Psi\rangle = |\psi\rangle_1 \otimes |\psi\rangle_2 = (\alpha_1 |0\rangle + \beta_1 |1\rangle) \otimes (\alpha_2 |0\rangle + \beta_2 |1\rangle).$$

Expanding this yields:

$$|\Psi\rangle = \alpha_1\alpha_2 |00\rangle + \alpha_1\beta_2 |01\rangle + \beta_1\alpha_2 |10\rangle + \beta_1\beta_2 |11\rangle.$$

***D7. Superposition and Entanglement:*** Superposition allows a quantum state to exist as a combination of basis states. *Entanglement*, a uniquely quantum phenomenon, occurs when qubits are so strongly correlated that the state of one qubit cannot be described independently of the other. For example, the entangled state:

$$|\Psi\rangle = \frac{1}{\sqrt{2}}(|00\rangle + |11\rangle),$$

cannot be written as a product of individual qubit states.

### *A. Quantum Gates*

Quantum gates are the building blocks of quantum circuits, analogous to classical logic gates. These gates operate on qubits and perform unitary transformations, preserving the normalization of the quantum state. Below, we explain some commonly used quantum gates, including their matrix representations and functions.

***E1. Pauli Gates:*** The *Pauli gates* represent single-qubit operations corresponding to rotations around the $X$, $Y$, and $Z$ axes of the Bloch Sphere.

- **Pauli-X Gate (NOT Gate)**: Flips the state of the qubit, mapping $|0\rangle$ to $|1\rangle$ and vice versa.

$$X = \begin{pmatrix} 0 & 1 \\ 1 & 0 \end{pmatrix}.$$

- **Pauli-Y Gate**: Represents a rotation around the $Y$-axis of the Bloch Sphere.

$$Y = \begin{pmatrix} 0 & -i \\ i & 0 \end{pmatrix}.$$

- **Pauli-Z Gate**: Represents a rotation around the $Z$-axis of the Bloch Sphere. It flips the phase of the $|1\rangle$ state.

$$Z = \begin{pmatrix} 1 & 0 \\ 0 & -1 \end{pmatrix}.$$

***E2. Hadamard Gate:*** The *Hadamard gate* creates superposition states by mapping $|0\rangle$ and $|1\rangle$ to equal superpositions of these states:

$$H = \frac{1}{\sqrt{2}} \begin{pmatrix} 1 & 1 \\ 1 & -1 \end{pmatrix}.$$

When applied to $|0\rangle$, the Hadamard gate produces:

$$H|0\rangle = \frac{1}{\sqrt{2}}(|0\rangle + |1\rangle).$$

***E3. Rotation Gates:*** Rotation gates perform rotations of a qubit state around a specific axis. These gates are parameterized by an angle $\theta$:

$$R_x(\theta) = \cos\left(\frac{\theta}{2}\right) I - i \sin\left(\frac{\theta}{2}\right) X = \\ = \begin{pmatrix} \cos\left(\frac{\theta}{2}\right) & -i\sin\left(\frac{\theta}{2}\right) \\ -i\sin\left(\frac{\theta}{2}\right) & \cos\left(\frac{\theta}{2}\right) \end{pmatrix}.$$

$$R_y(\theta) = \cos\left(\frac{\theta}{2}\right) I - i \sin\left(\frac{\theta}{2}\right) Y \\ = \begin{pmatrix} \cos\left(\frac{\theta}{2}\right) & -\sin\left(\frac{\theta}{2}\right) \\ \sin\left(\frac{\theta}{2}\right) & \cos\left(\frac{\theta}{2}\right) \end{pmatrix}.$$

$$R_z(\theta) = \cos\left(\frac{\theta}{2}\right) I - i \sin\left(\frac{\theta}{2}\right) Z \\ = \begin{pmatrix} e^{-i\theta/2} & 0 \\ 0 & e^{i\theta/2} \end{pmatrix}.$$

***E4. Controlled-NOT (CNOT) Gate:*** The *CNOT gate* is a two-qubit gate that flips the state of the target qubit if the control qubit is in the $|1\rangle$ state. Its matrix representation is:

$$\text{CNOT} = \begin{pmatrix} 1 & 0 & 0 & 0 \\ 0 & 1 & 0 & 0 \\ 0 & 0 & 0 & 1 \\ 0 & 0 & 1 & 0 \end{pmatrix}.$$

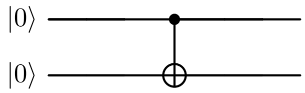


Fig. 14. The controlled-NOT (CNOT) gate. The filled circle denotes the control qubit and the circled plus denotes the target qubit.

In the computational basis:

- $\text{CNOT}|00\rangle = |00\rangle$,
- $\text{CNOT}|01\rangle = |01\rangle$,
- $\text{CNOT}|10\rangle = |11\rangle$,
- $\text{CNOT}|11\rangle = |10\rangle$.

***E5. Universal Gate Set:*** Any quantum circuit can be constructed using a combination of the following gates:

- Single-qubit gates (e.g., $H$, $R_x$, $R_y$, $R_z$),
- Two-qubit entangling gates (e.g., CNOT).

These gates form a *universal gate set*, capable of approximating any unitary operation on $n$-qubit systems.

ctionQuantum Data Encoding

In quantum computing, data from classical systems must be encoded into quantum states. This section discusses two encoding schemes used in our work: *Amplitude Encoding* and *Variational Encoding*. We demonstrate each encoding with an example. Note: the basics of quantum computing are covered in Appendix A.

*B. Amplitude Encoding*

Amplitude encoding maps a classical data vector into the amplitudes of a quantum state. The visual representation of the quantum circuit for amplitude encoding is provided in Fig. 15. Given a normalized classical vector $\mathbf{x} = [x_0, x_1, \ldots, x_{N-1}]$, it is encoded as a quantum state:

$$|\Psi\rangle = \sum_{i=0}^{N-1} \alpha_i |i\rangle, \quad \text{where } \alpha_i = \frac{x_i}{\|\mathbf{x}\|}, \ \|\mathbf{x}\| = \sqrt{\sum_{i=0}^{N-1} x_i^2}.$$

This encoding reduces the number of qubits needed for large datasets, as an $N$-dimensional vector requires only $\log_2(N)$ qubits.

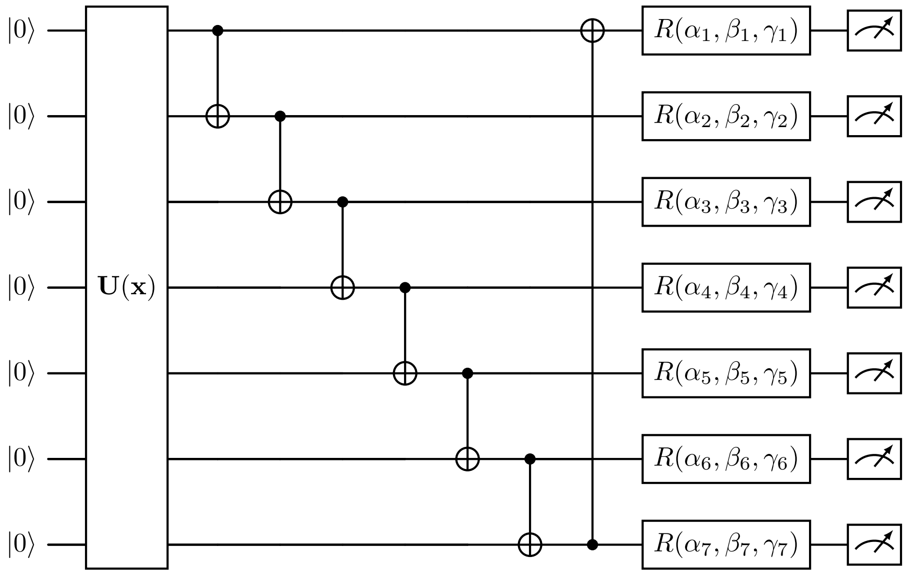


Fig. 15. Variational quantum circuit architecture for the classifier with amplitude encoding. The initial VQC block encodes the vector produced by the graph convolution operation. An $N$-dimensional vector is transformed into a quantum state using $\log_2(N)$ qubits. In this study, $N = 10$. The $U(\mathbf{x})$ function represents the quantum routine for amplitude encoding. Trainable parameters, denoted by $\alpha_i$, $\beta_i$, and $\gamma_i$, are optimized during training.

*a) Example:* Consider the classical data array $\mathbf{x} = [3, 8, 0, 4]$. First, normalize the vector to ensure it sums to unity:

$$\|\mathbf{x}\| = \sqrt{3^2 + 8^2 + 0^2 + 4^2} = \sqrt{9 + 64 + 0 + 16} = \sqrt{89}.$$

The normalized vector is:

$$\tilde{\mathbf{x}} = \left[\frac{3}{\sqrt{89}}, \frac{8}{\sqrt{89}}, \frac{0}{\sqrt{89}}, \frac{4}{\sqrt{89}}\right].$$

Using amplitude encoding, this vector is represented as a quantum state on 2 qubits (since $N = 4$):

$$|\Psi\rangle = \frac{3}{\sqrt{89}}|00\rangle + \frac{8}{\sqrt{89}}|01\rangle + \frac{0}{\sqrt{89}}|10\rangle + \frac{4}{\sqrt{89}}|11\rangle.$$

This process efficiently encodes a classical vector into a quantum state, reducing the dimensionality from $N$ to $\log_2(N)$ qubits.

*C. Variational Encoding*

Variational encoding maps classical data into quantum states by using the values in the data array as rotation angles for quantum gates, see Fig. 16 for the visual representation of the quantum circuit. For a given classical vector $\mathbf{x} = [x_1, x_2, \ldots, x_n]$, the encoding uses parameterized single-qubit gates, such as $R_y$ and $R_z$, to prepare the quantum state:

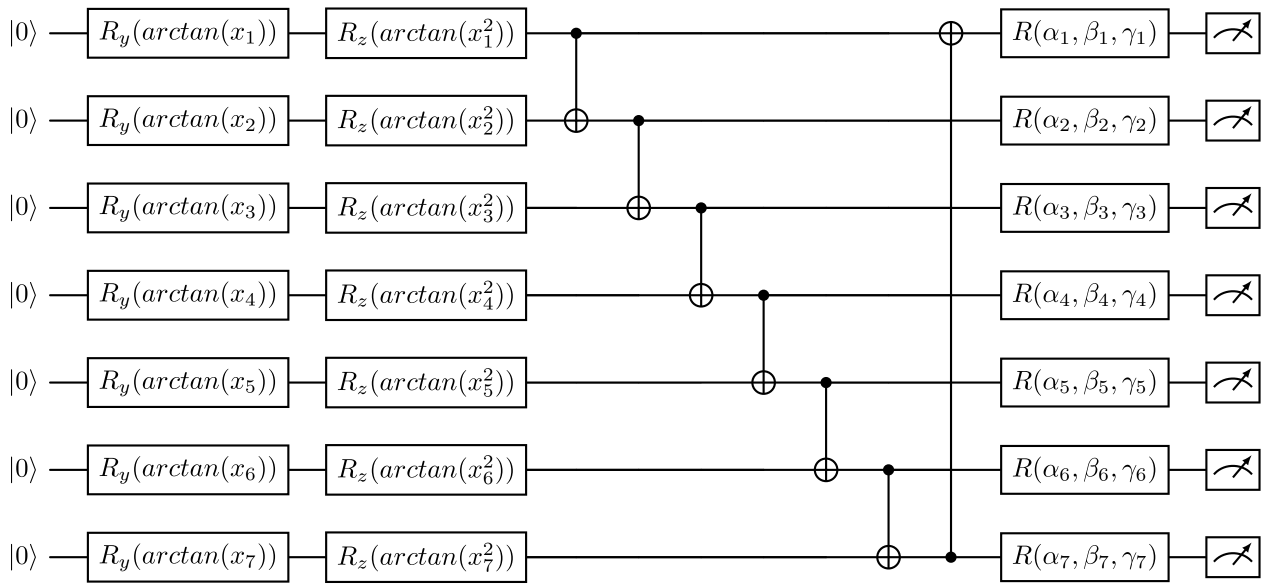


Fig. 16. Variational quantum circuit block with variational encoding. This encoding block employs $R_y$ and $R_z$ rotation gates, where the rotation angles are parameterized by $\arctan(x_i)$ and $\arctan(x_i^2)$ for each qubit. The parameters $\alpha_i$, $\beta_i$, and $\gamma_i$ are trainable and optimized during the learning process.

$$U(\mathbf{x}) = R_z(\arctan(x_1^2))R_y(\arctan(x_1))\otimes \cdots \otimes R_z(\arctan(x_n^2))R_y(\arctan(x_n)).$$

*a) Example:* Consider the same data array $\mathbf{x} = [3, 8, 0, 4]$. The variational encoding maps the elements into rotation angles for each qubit. For the $i$th element $x_i$, the angles are calculated as:

$$\theta_y = \arctan(x_i), \quad \theta_z = \arctan(x_i^2).$$

The angles (in radians) for each element are:

$$x_0 = 3: \quad \theta_y = \arctan(3), \theta_z = \arctan(3^2),$$
$$x_1 = 8: \quad \theta_y = \arctan(8), \theta_z = \arctan(8^2),$$
$$x_2 = 0: \quad \theta_y = \arctan(0), \theta_z = \arctan(0^2),$$
$$x_3 = 4: \quad \theta_y = \arctan(4), \theta_z = \arctan(4^2).$$

Each qubit undergoes rotations based on these angles using $R_y$ and $R_z$ gates. For instance, the first qubit is encoded as:

$$R_z\left(\arctan{(3)}^2\right) R_y\left(\arctan{(3)}\right)|0\rangle.$$

After applying variational encoding to all qubits, the quantum state reflects the classical data as rotation parameters.

### Appendix B
### Vina Distance Optimization Terms

In our study, the Vina distance optimization terms incorporate the same weighted scoring terms as used in AutoDock Vina [33]. These terms are composed of five conformation-dependent terms and one conformation-independent term. The five conformation-dependent terms represent intermolecular contributions, including three terms for steric interactions, one term for hydrophobic bonding, and one for hydrogen bonding. The detailed setup of Vina terms is available in Appendix B The single conformation-independent term accounts for flexibility, defined by the number of active rotatable bonds $N_{\text{rot}}$ between heavy atoms of the ligand. $N_{\text{rot}}$ can be calculated using RDKit [38]:

```
from rdkit import Chem

mol = Chem.MolFromMol2File('ligand.mol2')
N_rot = Chem.rdMolDescriptors.\
        CalcNumRotatableBonds(mol)
print("(N_rot):", N_rot) # e.g. 9, 5, etc.
```

The six Vina terms are mathematically defined as follows [33]:

$$\text{Gauss}_1(t_i, t_j, r_{ij}) = w_1 e^{-(d_{ij}/0.5)^2} \tag{1}$$

$$\text{Gauss}_2(t_i, t_j, r_{ij}) = w_2 e^{-((d_{ij}-3)/2)^2} \tag{2}$$

$$\text{Repulsion}(t_i, t_j, r_{ij}) = \begin{cases} w_3 d_{ij}^2 & \text{if } d_{ij} < 0 \\ 0 & \text{if } d_{ij} \geq 0 \end{cases} \tag{3}$$

$$\text{Hydrophobic}(t_i, t_j, r_{ij}) = \begin{cases} w_4 & \text{if } d_{ij} \leq 0.5 \\ w_4(1.5 - d_{ij}) & \text{if } 0.5 < d_{ij} < 1.5 \\ 0 & \text{if } d_{ij} \geq 1.5 \end{cases} \tag{4}$$

$$\text{HBonding}(t_i, t_j, r_{ij}) = \begin{cases} w_5 & \text{if } d_{ij} \leq -0.7 \\ w_5 \left(\frac{d_{ij}}{-0.7}\right) & \text{if } -0.7 < d_{ij} < 0 \\ 0 & \text{if } d_{ij} \geq 0 \end{cases} \tag{5}$$

$$\text{Flexibility} = w_6 N_{\text{rot}} \tag{6}$$

where $d_{ij} = r_{ij} - R_{t_i} - R_{t_j}$ represents the surface distance between atoms $i$ and $j$, excluding hydrogen atoms. Here, $t_i$ and $t_j$ are the atom types of atoms $i$ and $j$, respectively. $R_{t_i}$ and $R_{t_j}$ are the Van der Waals radii of these atom types, and $r_{ij}$ is their interatomic distance, with a cutoff at $r_{ij} = 8$ Å. All distances are Euclidean, and all units are in Å. In our case study, the dataset contained only 10 distinct types of atoms, and we employed Bondi van der Waals radii [39]:

```
vdw_radii = {
    8: 1.52,  # Oxygen
    7: 1.55,  # Nitrogen
    6: 1.7,   # Carbon
    16: 1.8,  # Sulphur
    35: 1.85, # Bromine
    15: 1.8,  # Phosphorus
    17: 1.75, # Cholorine
    9: 1.47,  # Fluorine
    53: 1.98, # Iodine
}
```

The weights $w_1$, $w_2$, $w_3$, $w_4$, $w_5$, and $w_6$ are taken from AutoDock Vina, with values of -0.0356, -0.00516, 0.840, -0.0351, -0.587, and 0.0585, respectively.
